# SARS-CoV-2 and SARS-CoV Spike-RBD Structure and Receptor Binding Comparison and Potential Implications on Neutralizing Antibody and Vaccine Development

**DOI:** 10.1101/2020.02.16.951723

**Authors:** Chunyun Sun, Long Chen, Ji Yang, Chunxia Luo, Yanjing Zhang, Jing Li, Jiahui Yang, Jie Zhang, Liangzhi Xie

## Abstract

SARS-CoV-2 and SARS-CoV share a common human receptor ACE2. Protein-protein interaction structure modeling indicates that spike-RBD of the two viruses also has similar overall binding conformation and binding free energy to ACE2. *In vitro* assays using recombinant ACE2 proteins and ACE2 expressing cells confirmed the two coronaviruses’ similar binding affinities to ACE2. The above studies provide experimental supporting evidences and possible explanation for the high transmissibility observed in the SARS-CoV-2 outbreak. Potent ACE2-blocking SARS-CoV neutralizing antibodies showed limited cross-binding and neutralizing activities to SARS-CoV-2. ACE2-non-blocking SARS-CoV RBD antibodies, though with weaker neutralizing activities against SARS-CoV, showed positive cross-neutralizing activities to SARS-CoV-2 with an unknown mechanism. These findings suggest a trade-off between the efficacy and spectrum for therapeutic antibodies to different coronaviruses, and hence highlight the possibilities and challenges in developing broadly protecting antibodies and vaccines against SARS-CoV-2 and its future mutants.

## Introduction

In December 2019, outbreak of SARS-CoV-2 infection has brought back the attention of pathogenic coronavirus to the spotlight^1-5^. SARS-CoV-2 is spreading rapidly, causing severe COVID-19 symptoms and life-threatening diseases in some infected patients^6^. Numbers of infected cases reached over 60,000 in less than 3 months^7^. Various estimates and analyses suggest SARS-CoV-2 and Severe Acute Respiratory Syndrome coronavirus (SARS-CoV) may have similar transmissibility with an estimated reproductive number (R_0_) of 3.77 (2.23-4.82) for SARS-CoV-2 and between 2.9-3.3 for SARS-CoV^8^.

SARS-CoV-2 and SARS-CoV share a common host-cell receptor protein, angiotensin converting enzyme 2 (ACE2), expressed on epithelial cells in the respiratory track system and various human organs, such as the lung^9,10^. Receptor recognition by coronaviruses is the first and essential step for infecting host cells^11,12^. An envelope-anchored trimeric spike (S) protein mediates the binding of the two coronaviruses to human ACE2^13,14^, and is cleaved by the host protease into two separate polypeptides as S1, which contains the receptor binding domain (RBD), and S2, which is responsible for the homotrimer formation and mediates fusion of the virion with cellular membranes^15^. The receptor binding motif (RBM) in RBD is responsible for direct binding to ACE2 and its binding affinity may directly affects the virus infectivity and transmissibility^16,17^. The amino acid sequence homology between SARS-CoV and SARS-CoV-2 approximates to 75% for the spike proteins, and are 73.7% and 50.0% for RBDs and RBMs, respectively.

The prominent sequence differences between the crucial RBMs of SARS-CoV-2 and SARS-CoV raise a critical question of whether the binding affinity of SARS-CoV-2 spike protein to human ACE2 is comparable to that of SARS-CoV. A recent study by computational modeling suggested that SARS-CoV-2 has a lower binding affinity to human ACE2, as a result of the loss of one hydrogen bond interactions^18^. However, another publication using structure analysis suggested similar binding affinities to SARS-CoV^9^. By using biolayer interferometry binding assay, Tian et al.^9^ measured the SARS-CoV-2 RBD’s binding affinity to human ACE2 protein to be 15.2 nM, which is comparable to previously published affinity data for SARS-CoV spike protein^19^. The above comparison of spike proteins’ binding affinities to human ACE2 between SARS-CoV-2 and SARS-CoV were inconclusive or indirect, hence, a direct head-to-head comparison is desired for the understanding of the infectivity and transmissibility of SARS-CoV-2 virus.

Neutralizing antibody (nAb) is expected to be one of the most promising treatments against coronavirus infection among the existing therapeutic options^20,21^. Several nAbs targeting SARS-CoV exhibit significant *in vivo* antivirus activities by reducing virus titers in lung tissues of animal models^22-26^. However, coronavirus is a single-stranded RNA virus prone to rapid mutations during transmission, nAbs without cross-reactivity to a broad spectrum of viral mutants could lead to treatment failure^10,27-29^, therefore, highly potent and cross-protective nAbs and prophylactic vaccines against SARS-CoV-2 are in urgent needs.

Here we investigated the optimized complex structure conformations between the RBDs of SARS-CoV-2 (Wuhan/IVDC-HB-01/2019, GISAID accession ID: EPI_ISL_402119) and SARS-CoV (CUHK-W1, GenBank accession ID: AY278554) with ACE2 by computational modeling and binding free energy analysis. We also used the recombinant S1 proteins of the two viruses to compare their binding curves to both recombinant ACE2 protein and ACE2 expressing cells. The above studies confirm that both SARS-CoV-2 and SARS-CoV have similar binding affinities to the human receptor ACE2.

Due to the relatively low homology of the spike RBDs between SARS-CoV-2 and SARS-CoV, it is of significant interest to investigate whether SARS-CoV neutralizing antibodies possess cross-reactivity to SARS-CoV-2. SARS-CoV polyclonal antibodies and ACE2 blocking and non-blocking nAbs were tested with SARS-CoV-2 pseudovirus (PSV) for cross neutralizing activities.

## Results

### SARS-CoV-2 RBD-ACE2 complex structure modeling reveals almost identical structure configuration and affinity to SARS-CoV

The structure model for SARS-CoVRBD-ACE2 complex was optimized based on the complex’s crystal structure (PDB ID: 2AJF) (http://www.rcsb.org/). The structure model for SARS-CoV-2 RBD-ACE2 complex was constructed by homology modeling using the optimized SARS-CoV RBD-ACE2 complex crystal structure as a template, based on a 73.7% amino acid sequence homology between the two viruses. Root-mean-square deviation (RMSD) of Cα atoms between the two complex structures is 0.703 Å, indicating an almost identical structural conformation (**Figure1B**). There are subtle differences in the RBD-ACE2 interfaces between SARS-CoV-2 and SARS-CoV at the loop 469-470 (numbered according to SARS-CoV RBD), arising from a one-residue insertion after residue 469 (**Figure 1A**).

**Figure 1.**
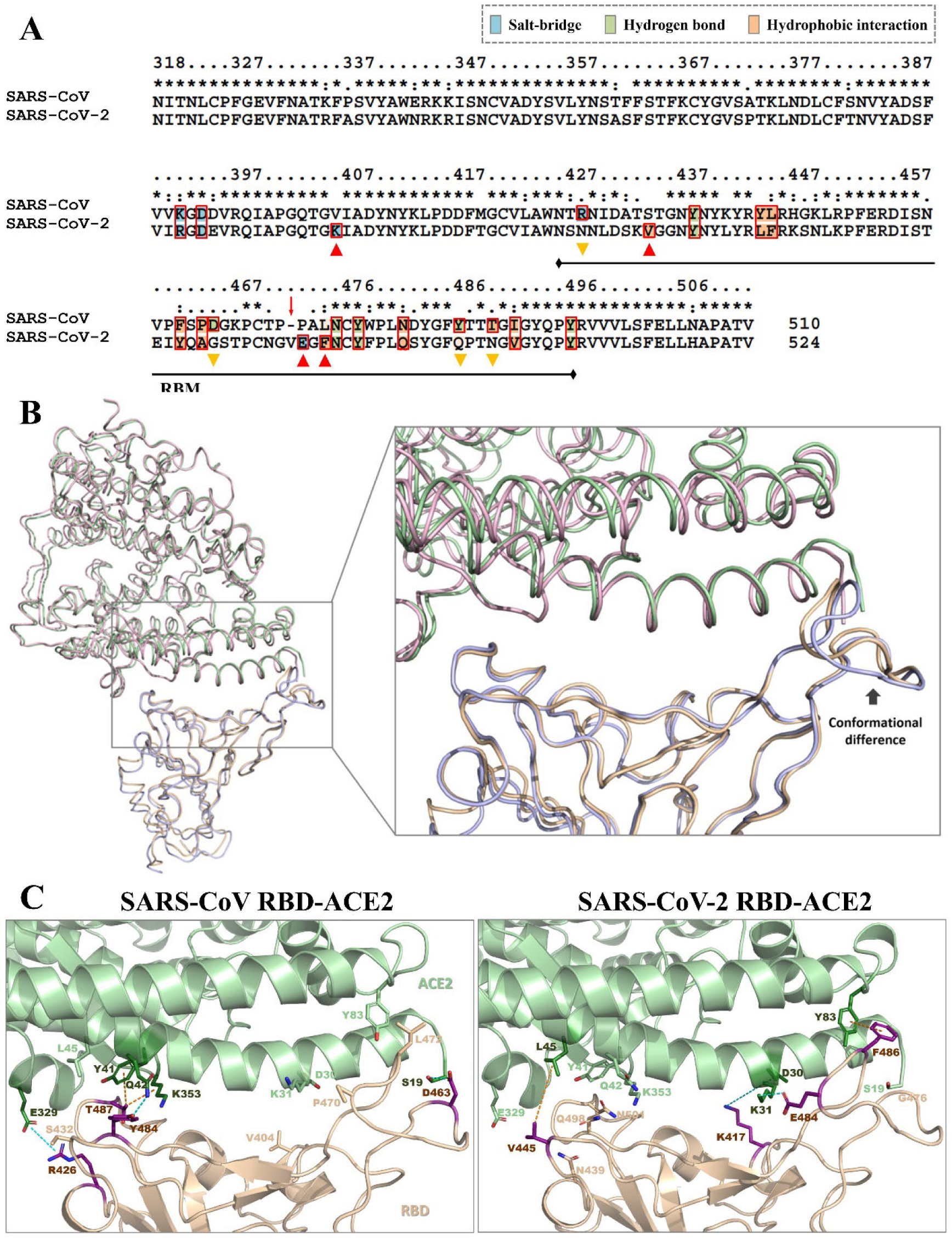
Sequence analysis and structure modeling of SARS-CoV-2 RBD and SARS-CoV RBD and their interactions with ACE2. **A.** RBD sequence alignment of SARS-CoV and SARS-CoV-2, highlighting the predominant residues that contribute to the interactions with ACE2. The distinct interactions of RBD and ACE2 for the two viruses are indicated by the down-pointing orange triangles and up-pointing red triangles, respectively. RBM residues are underlined. The one-residue insertion is indicated by the red arrow. Asterisks indicate positions of fully conserved residues. Colons indicate positions of strictly conserved residues. Periods indicate positions of weakly conserved residues. **B.** Conformational comparison between the RBD-ACE2 complex structures for SARS-CoV-2 and SARS-CoV. The RBD and ACE2 structures in the SARS-CoV-2 RBD-ACE2 complex model are shown as orange and pink tubes, respectively. The RBD and ACE2 structures in the optimized SARS-CoV RBD-ACE2 complex structure are shown as blue and green tubes, respectively. The location of noticeable subtle conformational difference is indicated by an arrow. **C.** Distinct interaction patterns in the SARS-CoV-2 and SARS-CoV RBD-ACE2 interfaces. Structures of RBD and ACE2 are shown as cartoon in pink and green colors, respectively. The side chains of the residues in both protein components, representing their unique interactions, are shown as sticks. Polar interactions (salt-bridge and hydrogen bond) are shown as blue dash line. Non-polar interactions (π-stack, π-anion, and hydrophobic interactions) are shown as orange dash line.

On the other hand, the interaction patterns in both complex structures’ interfaces are somewhat different (**Figure 1A and Table S2**). Four residues in SARS-CoV-2 RBD (i.e. N439, G476, Q498, and N501) lost their interactions with ACE2 as compared with the corresponding residues in SARS-CoV RBD (i.e. R426, D463, Y484, and T487), four replacing residues in SARS-CoV-2 RBD (i.e. K417, V445, E484, and F486) form new interactions with ACE2, which are absent in the SARS-CoV RBD (corresponding to V404, S432, P470, and L472) (**Figure 1A&C**). It is noteworthy that although a strong salt-bridge presented between R426 of SARS-CoV RBD and E329 of ACE2, is missing in the interaction involving SARS-CoV-2 RBD. However, a new strong salt-bridge interaction between E484 of SARS-CoV-2 RBD and K31 of ACE2 compensates for that loss.

The RBD-ACE2 binding free energies of SARS-CoV and SARS-CoV-2 are estimated to be −40.42 and −44.96 REU (Rosetta energy unit), respectively (**Table S1**), by Rosetta Interface Analyzer^30^. The insignificant difference in binding free energies suggests that SARS-CoV-2 and SARS-CoV viruses have similar binding affinities to human ACE2.

### Recombinant S1 protein of SARS-CoV-2 showed similar binding to human ACE2 in both protein and cellular forms, as compared with SARS-CoV

The RBD containing S1 protein, resulted from cleavage of the spike protein on the virus membrane is responsible for the binding of the virus to human receptor on cell membrane, which is important for viral infectivity^31^. Binding curves of recombinant S1 proteins of SARS-CoV-2 and SARS-CoV to human ACE2 were measured by ELISA. Results confirmed that both viruses have similar S1-ACE2 protein-protein binding curves and EC_50_ (**Figure 2A**). Binding curves of S1 protein to ACE2 expressing 293T cells was further assessed using FACS. Again, similar binding curves and EC_50_ values were obtained for the two viruses (**Figure 2B**). The above experimental results are consistent with our structure modeling analysis, which indicates that SARS-CoV-2 virus likely infects human cells through similar mechanisms as SARS-CoV virus by binding to human ACE2 with comparable affinities, and hence may possess similar transmissibility.

**Figure 2.**
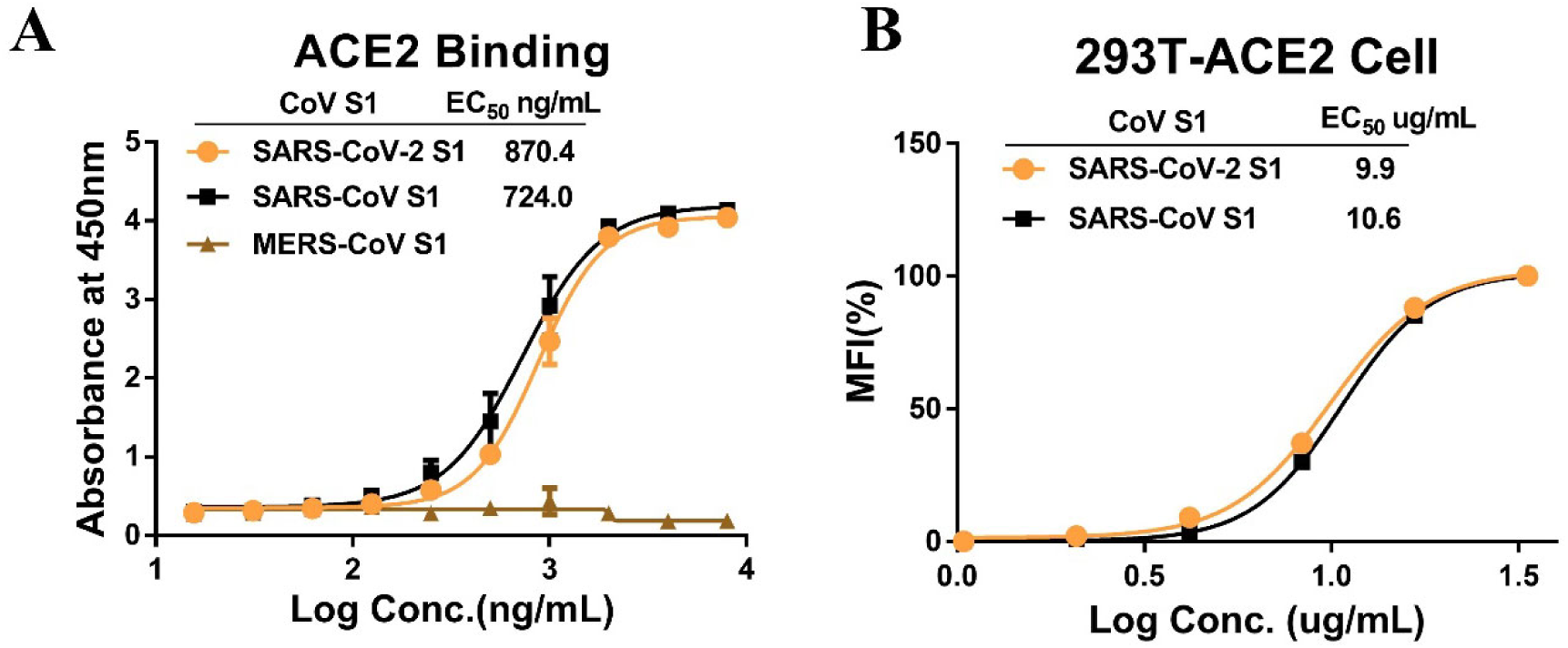
Measurements of SARS-CoV-2 and SARS-CoV S1 binding to ACE2. A. Serial diluted recombinant S1 proteins of SARS-CoV-2, SARS-CoV and MERS-CoV were coated on 96 well plates, incubated with the recombinant Fc-tagged ACE2 (ACE2-Fc) for binding evaluation. B. Recombinant S1 proteins of SARS-CoV-2 and SARS-CoV were incubated with 293T-ACE2 cells and subjected to FACS evaluation for binding.

**Figure 3.**
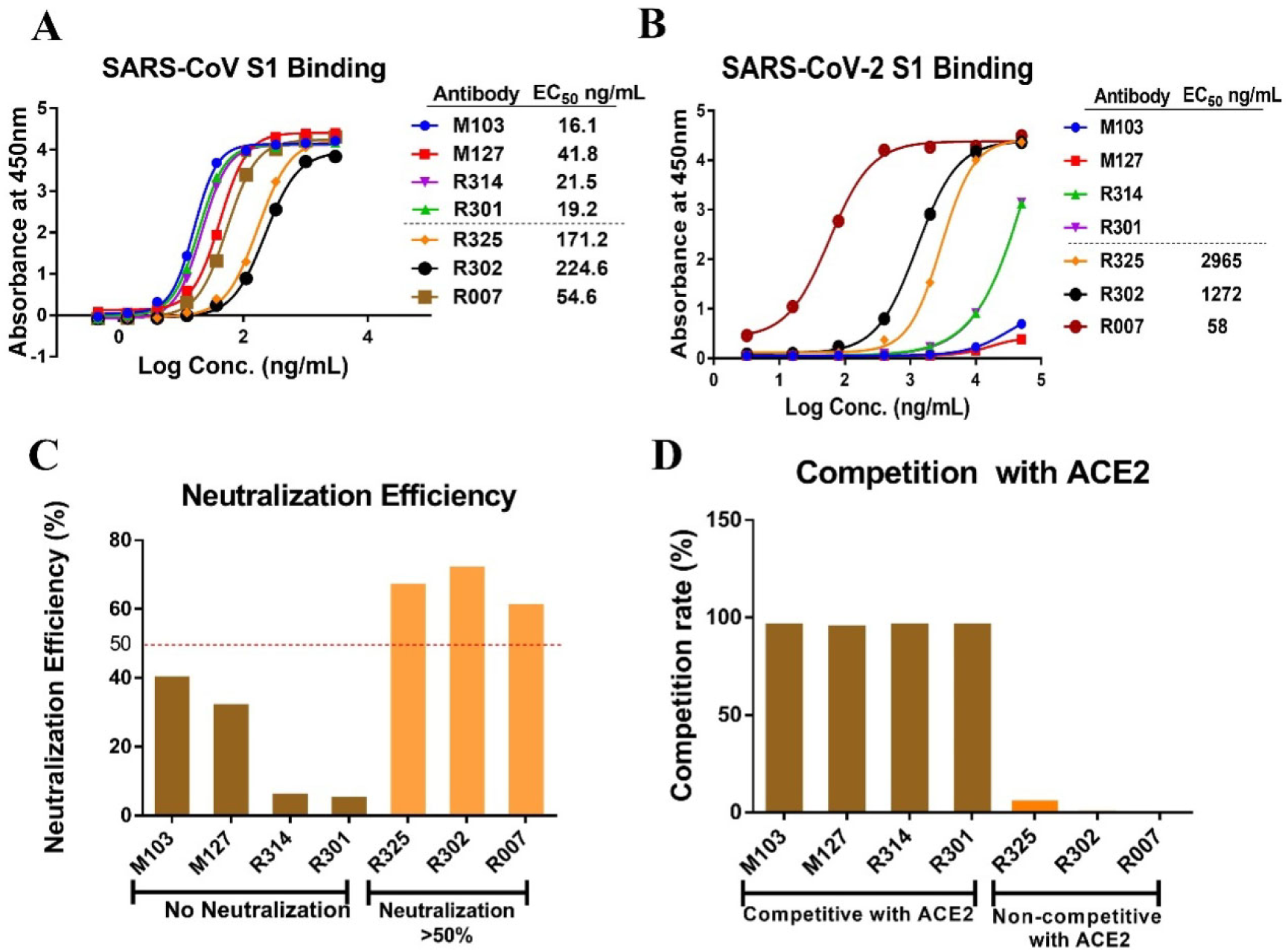
Cross-reactivity and neutralization efficiency of SARS nAbs against SARS-CoV-2. A. Binding of SARS nAbs to SARS-CoV S1 protein were tested by ELISA. Recombinant S1 protein of SARS-CoV were coated on plates, serial diluted nAbs were added for binding to recombinant S1 protein. B. Binding of SARS nAbs to SARS-CoV-2 S1 protein were tested by ELSIA. Recombinant S1 protein of SARS-CoV-2 were coated on plates, serial diluted nAbs were added for binding to recombinant S1 protein. C. Neutralization of SARS-CoV nAbs against SARS-CoV-2 PSV. D. Antibody competition with SARS-CoV RBD binding to ACE2. Recombinant SARS-CoV RBD protein was coated on plates, nAbs and recombinant ACE2 were then added for RBD binding competition measurements.

### ACE2-non-blocking SARS-CoV nAbs showed modest S1-protein binding and pseudovirus neutralizing activities to SARS-CoV-2

Coronavirus’ mutation and emergence of escape mutants to neutralizing monoclonal antibodies and vaccines are a major concern. SARS-CoV-2 and SARS-CoV are two different coronaviruses with modest level of sequence homology in their spike proteins. Understanding whether antibodies raised from SARS-CoV spike protein immunization have cross-reactivity to the new SARS-CoV-2 will offer important insights and guidance to therapeutic antibody and prophylactic vaccine development.

SARS-CoV nAbs were generated by immunizing mouse or rabbit with SARS-CoV S1 or RBD protein. Two SARS-CoV S-protein rabbit polyclonal antibodies **(Table 1)** and four monoclonal antibodies **(Table 2)** were analyzed for cross-reactivity to SARS-CoV-2 S1 protein and cross-neutralizing activities to SARS-CoV-2 PSV. As expected, the polyclonal antibodies showed weaker binding to SARS-CoV-2 S1 protein (EC_50_>100ng/mL) as compared to SARS-CoV S1 protein (EC_50_<25ng/mL). Neutralizing activities against SARS-CoV-2 PSV were lower by more than two-orders of magnitudes than SARS-CoV PSV **(Table 1)**, presenting a pessimistic forecast for the probability to identify highly potent and cross-reactive nAbs to SARS-CoV-2 from SARS-CoV antibodies or antibody libraries.

**Table 1.** Polyclonal nAbs characteristics.

| Antibody<br>(Polyclonal) | Pseudovirus Neutralization |  | S1 Protein Binding |  |
| --- | --- | --- | --- | --- |
|  | IC<sub>50</sub> (μg/mL) |  | EC<sub>50</sub> (ng/mL) |  |
|  | SARS-CoV-2 | SARS-CoV | SARS-CoV-2 | SARS-CoV |
| RP01 | 137 | 1.1 | 130 | 21.5 |
| T52 | 50.4 | 0.08 | 113.6 | 7.9 |

RP01: polyclonal antibody from rabbit immunized with SARS-CoV S1 protein.
T51: polyclonal antibody from rabbit immunized with SARS-CoV RBD protein

**Table 2.** Monoclonal nAbs characteristics.

| Antibody |  | Pseudovirus Neutralization |  | S1 Binding Cross-reactivity |  | Affinity Constant |
| --- | --- | --- | --- | --- | --- | --- |
|  |  | Neutralization | Minimum dose | EC<sub>50</sub> (ng/mL) |  | K<sub>D</sub>(M) |
|  |  | at 100 µg/mL | 100%(µg/mL)<sup>#</sup> | SARS-CoV-2 | SARS-CoV | SARS-CoV S1 |
| Competitive with ACE2 | M103 | -* | 0.1 | -* | 16 | 8.7E-12 |
|  | M127 | -* | 0.1 | -* | 42 | 2.0E-10 |
|  | R314 | -* | 0.1 | -* | 21 | 5.0E-11 |
|  | R301 | -* | 0.1 | -* | 19 | 1.2E-11 |
| Not | R325 | 68% | 10 | 2965 | 171 | -* |
| Competitive | R302 | 73% | 10 | 1272 | 224 | -* |
| with ACE2 | R007 | 62% | >10 | 58 | 55 | -* |

<sup>#</sup>: minimum dose to reach 100% neutralization. \*: not feasible with similar assessment

SARS-CoV monoclonal nAbs with strong S1 binding (EC_50_<50ng/mL, K_D_≤2.0E-10 M) and potent ACE2-blocking activities exhibited potent neutralizing activities against SARS-CoV (minimum dose to reach 100% neutralization at ∼0.1μg/mL), but almost no cross-binding to SARS-CoV-2 S1 protein (EC_50_>15μg/mL) and no cross-neutralizing activities against SARS-CoV-2 PSV (<50% neutralizing activity at 100μg/mL) **(Figure3, Table 2, Figure S1 & S2)**.

Interestingly, three ACE2-non-blocking monoclonal antibodies showed modest binding activities to SARS-CoV S1 protein (EC_50_>100ng/mL) and neutralizing activities to the SARS-CoV PSV (minimum dose to reach 100% neutralization at ≥ 10 μg/mL). Although, these nAbs were less potent than the ACE2-blocking nAbs to SARS-CoV, but could evidently cross-bind to the SARS-CoV-2 S1 protein (EC_50_<3μg/mL) and cross-neutralize SARS-CoV-2 PSV (>50% neutralizing activity at 100μg/mL) **(Table 2)**.

### Significant micro-structure differences between SARS-CoV-2 and SARS-CoV RBDs exist in their RBM regions

As suggested by the SARS-CoV and SARS-CoV-2 RBD sequence alignment analysis (**Figure 1A**), the 50% RBM sequence homology is much less than the 73.7% RBD sequence homology. Structure similarity analysis confirms the presence of significant local structure differences in the RBM regions, while the rest of the RBD regions have significantly higher similarities (**Figure 4**). Moreover, three glycosylation sites in SARS-CoV RBD (i.e. N318, N330, and N357)^32^ was conserved, although one of the glycosylation sites, N357, may not be glycosylated in the corresponding residue of SARS-CoV-2 as predicted by NetNGlyc^33^.

**Figure 4.**
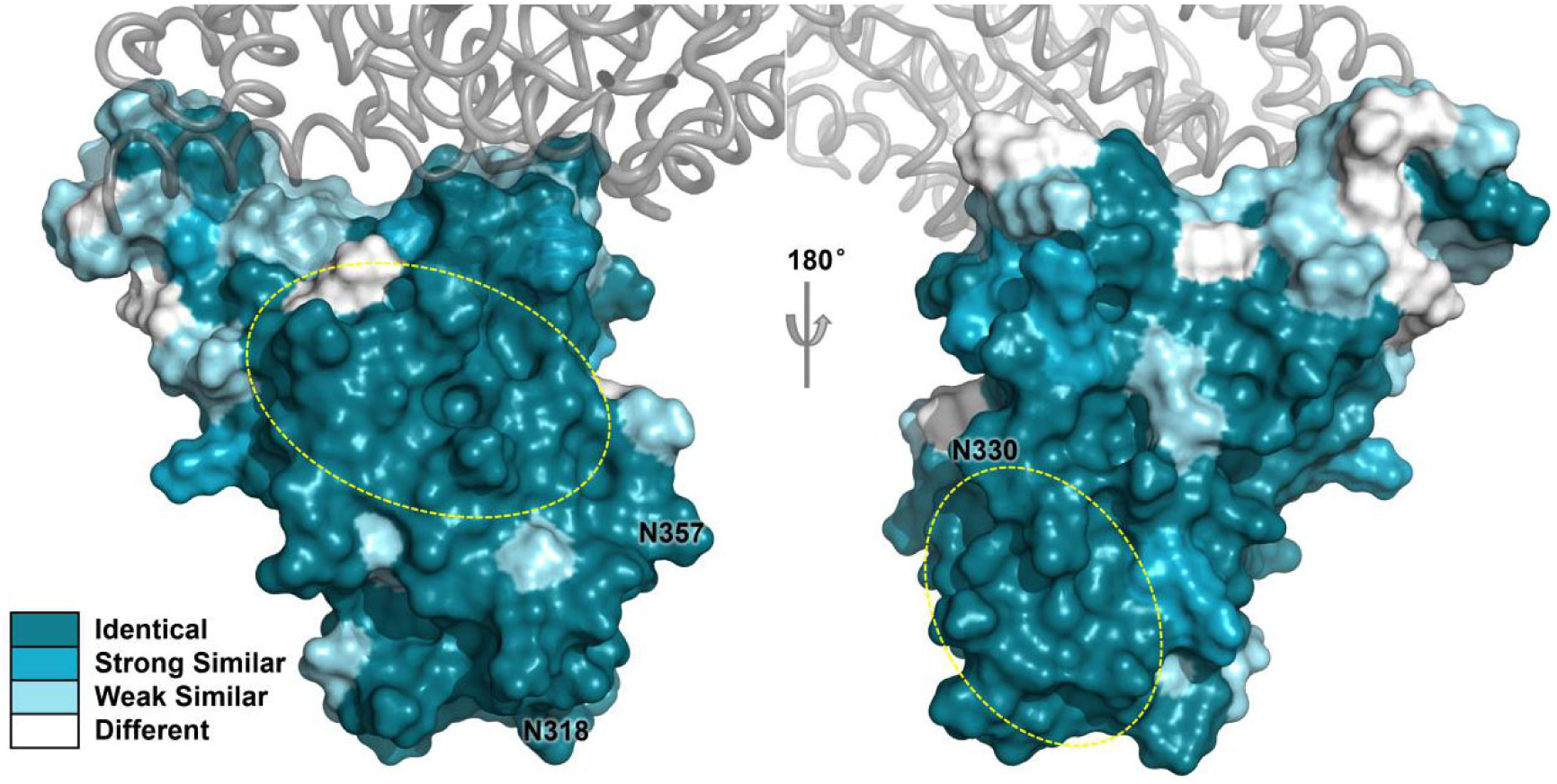
Structure similarity between SARS-CoV-2 RBD and SARS-CoV RBD. RBD is shown in a space-filled model with colored surface. ACE2 is shown as gray tube model. The three glycosylation sites in SARS-CoV are labeled. Note that N^357^ST in SARS-CoV is changed to N^370^SA in SARS-CoV-2, which is different from the NXS/T pattern required for glycosylation, and hence this site is more likely to be unglycosylated. The two possible cross-reactive regions are marked with yellow circles.

Taking the two viruses’ RBD structure similarities and glycosylation sites into account, we predict that there are two possible conserved regions in RBD where epitopes for cross-reactive neutralizing antibodies reside. These regions probably do not overlap with the ACE2 binding sites (**Figure 4**). We speculate that it is more likely to obtain cross-reactive and non-blocking neutralizing antibodies targeting the conserved regions outside the RBM region. The neutralization mechanism for these non-blocking but cross-reactive antibodies is likely unrelated to ACE2 blockage.

## Discussion

Coronavirus spike protein has been shown to be responsible for interacting with host cell receptors to initiate infection. Both SARS-CoV and SARS-CoV viruses use human ACE2 for cell entry^10^. Due to the nature of RNA virus’ rapid mutation rates, changes in the S-protein’s amino acid sequence, especially in the RBD’s receptor binding motif, could have significant impact on virus infectivity, pathogenicity, transmissibility, and cross-protection from previous coronavirus infection, as well as therapeutic antibody and prophylactic vaccine development. Hence, understanding the differences between SARS-CoV and SARS-CoV-2 and their implications may offer significant scientific and practical value.

Though it is difficult to assess the virus-host-cell interactions with real virion, we studied the new SARS-CoV-2 in head-to-head comparisons with SARS-CoV using multiple methods. Measurements of recombinant SARS-CoV-2 and SARS-CoV spike proteins to recombinant ACE2 protein and ACE2 expressing 293T cells confirmed the two coronaviruses’ similar binding affinities to human ACE2 (**Figure 2A&B**), which provided direct molecular based evidences to support and possibly explain the observation that the new SARS-CoV-2 coronavirus has similar transmissibility to SARS-CoV virus.

Since SARS-CoV-2 and SARS-CoV are both coronaviruses with over 70% sequence homology and share the same human receptor ACE2, analyzing SARS-CoV’s antibodies’ cross-reactivity to SARS-CoV-2 may provide useful information on whether neutralizing epitopes were conserved on the two coronaviruses. Two rabbit polyclonal antibodies produced with SARS-CoV S1 and RBD proteins had potent binding and neutralizing activities to SARS-CoV but only modest cross-binding and cross-neutralizing activities to the new SARS-CoV-2 virus. Four highly potent ACE2 blocking SARS-CoV monoclonal antibodies showed binding affinities to SARS-CoV S1 protein in the range of 0.2 nM to 8.7 pM, and very high neutralizing activities to SARS-CoV. As low as 0.2 ug/mL nAb can lead to 100% neutralization of SARS-CoV PSV. However, virtually no cross-binding or cross neutralizing activities against the novel SARS-CoV-2 virus were detected with the four ACE2 blocking monoclonal antibodies.

We then screened non-ACE2-blocking antibodies raised from SARS-CoV RBD for neutralizing activities. Three such monoclonal antibodies were identified. Although, binding affinities to SARS-CoV S1 protein was significantly lower with EC_50_ between 55 to 224 μg/mL as compared to 16 to 42 μg/mL EC_50_ for the four ACE2-blocking nAbs, significant cross-binding activities to SARS-CoV-2 S1 protein and modest cross-neutralizing activities against SARS-CoV-2 PSV were detected.

The observation that these three antibodies bind to and neutralize both SARS-CoV and SARS-CoV-2 without blocking ACE2 suggest the following. First, the epitope or epitopes for these three antibodies are conserved across the two significantly different coronaviruses. Hence it is possible but maybe challenging to identify antibodies with potent neutralizing activities to both SARS-CoV and SARS-CoV-2, and ideally to their mutant virus strains as well. Second, the epitope or epitopes for the three nAbs are likely located outside the RBM motif due to their non-ACE2 blocking features and the fact that homology in the RBM motif between the two viruses is significantly lower than the rest portion of the RBD.

An analysis on the mutations in RBD was also conducted. We analyzed 68 sequences of SARS-CoV-2 variants in GISAID and found 4 variants with mutations in RBD (one has N354D/D364Y, two have V367F, and one has F342L). Alignment of 111 SARS-CoV RBD sequences, collected by BLAST via NCBI website^34^, was used for ConSurf analysis^35^. As shown in **Figure 5**, the SARS-CoV RBM region is highly variable, making it more challenging to develop cross-reactive antibodies with broad spectrum of neutralizing activities against mutant strains. On the other hand, significant portions of RBD (marked in pink) outside the RBM motif are highly conserved. Neutralizing antibodies targeting epitopes in these regions could potentially have cross-protective activities against different mutant strains.

**Figure 5.**
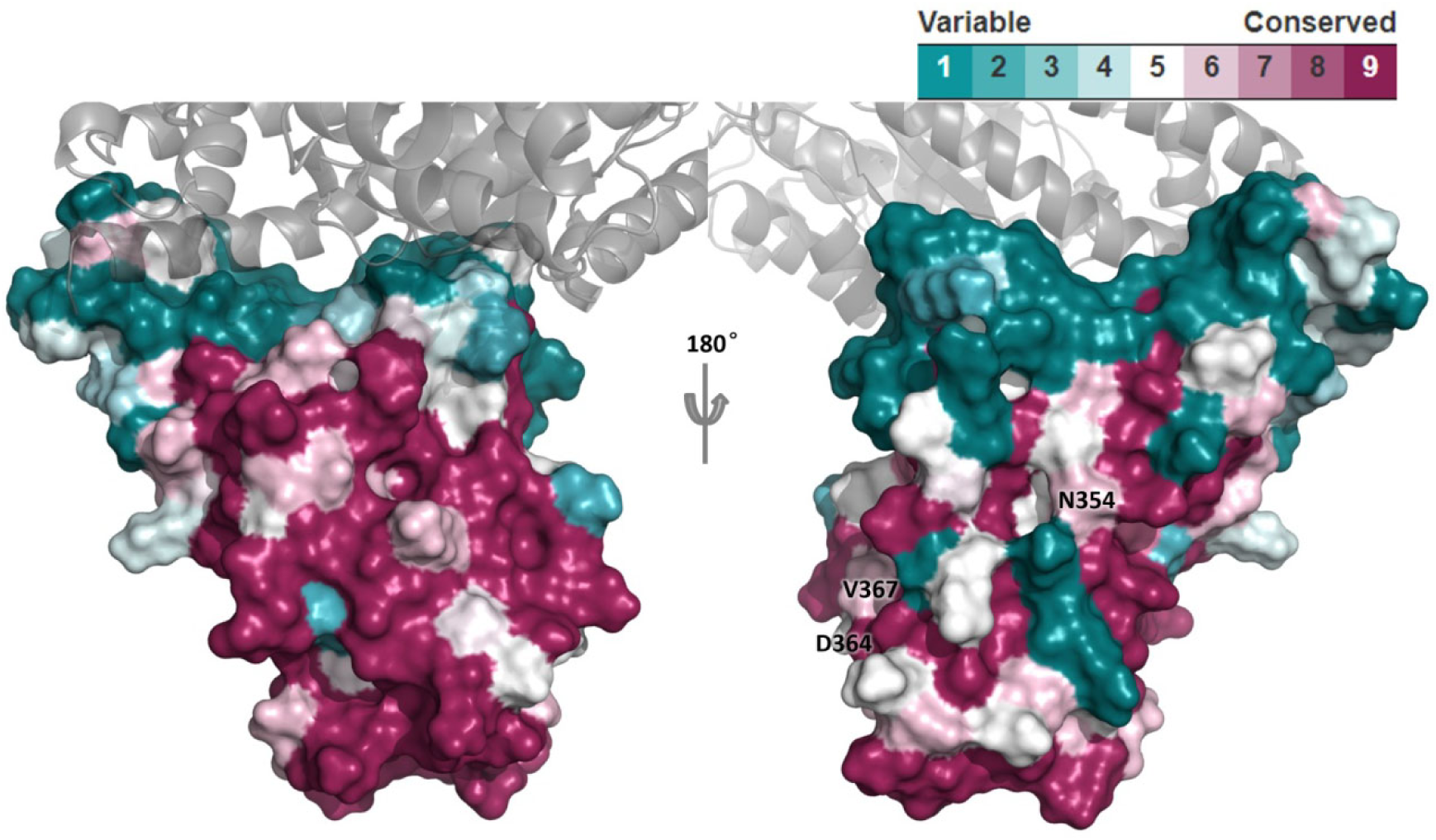
Structural conservation of SARS-CoV RBD. RBD is shown as colored surface. ACE2 is shown as gray cartoon. The three surface mutation sites (i.e. N354D, D364Y, and V367F) observed in SARS-CoV-2 RBD are labeled. Mutation F342L is buried and not shown here.

It is also noteworthy that the three non-blocking antibodies’ neutralizing activities are significantly lower than the four blocking antibodies. Due to the limited number of antibodies obtained, it is difficult to conclude whether there is a consistent pattern between ACE2 blockage and neutralizing potency, but one can speculate that neutralizing antibodies targeting conserved epitopes outside the RBM region may be cross-protective but may also be less potent due to the lack of ACE2 blocking activities. In a recent article^9^, a SARS-CoV RBD neutralizing antibody CR3022 with an epitope outside the RBM motif was also shown to be non-blocking but cross-binding to SARS-CoV-2 RBD with a relatively low K_D_ of 6.3 nM.

The neutralization mechanism of these non-blocking antibodies is not clear yet. It is known that antibodies targeting the S2 region could block S2 protein’s configuration change and hence interfere with virus entry into host cells. Whether these non-blocking RBD antibodies also interfere with S2 protein’s configuration change remains to be investigated.

In summary, the above findings suggest that SARS-CoV-2 and SARS-CoV bind to human ACE2 with similar affinities and hence may have similar transmissibility. Antibodies targeting the RBM regions may be more potent due to their ACE2 blocking activities but cross-protecting antibodies targeting the RBM regions may be more difficult to obtain because of the lower degree of sequence conservation. Antibodies to the more conserved regions outside the RBM motif may possess cross-protective neutralizing activities but may be less potent due to the lack of ACE2 blocking activities. Mechanism of neutralization for the non-blocking RBD antibodies remains to be investigated. Developing potent and cross-protective therapeutic antibodies and vaccines is possible but could be challenging.

## Materials and Methods

### Reagents, recombinant proteins and antibodies

Recombinant S1 proteins of SARS-CoV-2 (Cat: 40591-V08H), SARS-CoV (Cat: 40150-V08B1) and MERS-CoV (Cat:40069-V08H), recombinant RBD protein of SARS-CoV (Cat: 40150-V31B2), transfection reagent Sinofection (Cat: STF02), mammalian expression plasmids of full length S or RBD protein of SARS-CoV-2 (Cat: VG40589-UT, Wuhan/IVDC-HB-01/2019) and SARS-CoV (Cat: VG40150-G-N, CUHK-W1), ACE2 (Cat: HG10108-UT), polyclonal antibodies against SARS-CoV RP01 (Cat: 40150-RP01) and T52 (Cat: 40150-T52) were purchased from Sino Biological. Fetal bovine serum (FBS) (Cat: SA 112.02) were purchased from Lanzhou Minhai Bio-engineering. Hygromycin (Cat: V900372) were purchased from Sigma-Aldrich. SARS-CoV neutralizing antibodies were generated from mice (M103, M127) or rabbits (R314, R301, R325, R302, R258, R348) immunized with recombinant S1 protein of SARS-CoV. Luciferase assay system (Cat: E1501) was purchased from Promega.

### Cell lines

The human embryonic kidney 293T cell line (Cat:CRL-11268) used for pseudovirus (PSV) packaging were purchased from ATCC. 293T-ACE2 cells were established by transfection of ACE2 expression plasmid to 293T cells. Both 293T and 293T-ACE2 cells were grown in Dulbecco’s modified Eagle’s medium (DMEM) containing 10% (v/v) FBS.

### ELISA assay

Indicated proteins were coated on 96 well plates using CBS buffer over night at 4°C. BSA was used for blocking at room temperature for 1h. Indicated corresponding proteins or antibodies were then added and incubated at room temperature for 1h. After washing away the unbound proteins or antibodies, secondary antibody with HRP labeling were added and incubated for another hour before washed away. Developing buffer was added and incubated for 5-30 min, 1% H_2_SO_4_was added to stop the reaction and absorbance at 450nm was detected with a microplate reader.

### Flow cytometry

Indicated concentrations of S1 proteins of SARS-CoV-2 and SARS-CoV were incubated with 293T-ACE2 cells for 45 min. After washing away the unbound proteins, cells were incubated withPE labelled anti-his-tag antibody for 20 min and went through flow cytometer for detection of cellular binding. Flowjo and Graphpad softwares were used for data analysis.

### Octet

Recombinant SARS-CoV S1 protein were biotinated and loaded using SA sensor, indicated antibodies were added for real time association and dissociation analysis. Data Analysis Octet was used for data processing.

### Pseudovirus production in 293T adherent cells

6-8 hours before transfection, 293T cells were pre-plate on T75 flask in DMEM+10% FBS at 100,000 cells/cm^2^. 13μg of Luciferase-expressing HIV-1 lentiviral transfer genome (pWPXL-luc), 13μg of packaging plasmid (PSD) and 13μg of expression plasmid encoding either SARS-CoV-2-S protein (pCMV-whCoV-Spike) or SARS-S protein (pCMV-SARS-Spike) were co-transfected into pre-plated 293T cells using Sinofection transfection reagent according to the procedure recommended by manufacturer. Then the transfected cells incubated were incubated at 37°C and 5% CO_2_ overnight, followed by medium exchange with fresh DMEM plus 10% FBS. The supernatant containing pseudovirus was collected after 48-72 hours and filtered through a 0.45μm filter and stored at −80°C for longtime storage or 4°C for short time storage.

### Pseudovirus neutralization assay

293T-ACE2 cells were plated in 96-well plate at 1,0000 cells/well in 100 μ L DMEM+10% FBS. 60μL serially diluted antibody samples and 60μL of SARS-CoV-2 or SARS PSV were mixed and incubated at 4°C for 1 hour. Then 100μL/well of the antibody-PSV mixture was added onto the 293T/ACE2 cell wells and incubated for 37°C, 5% CO_2_. After 72 hours infection, luciferase luminescence (RLU) was detected using luciferase assay system according to the procedure recommended by manufacturer with a luminescence microplate reader. Antibodies inhibition% was calculated as following: 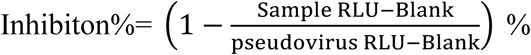. Antibodies neutralization titers were presented as 50% maximal inhibitory concentration (IC_50_).

### Sequence alignment

Protein sequence alignments were performed using Clustal Omega^36^.

### RBD-ACE2 complex structure modeling and analysis

The SARS-CoV RBD-ACE2 complex structure was remodeled based on PDB 2AJF to complete its missing loop using Discovery Studio (DS)^37^. This complex structure was then optimized by Rosetta Relax^38^. The SARS-CoV-2 RBD-ACE2 complex structure was constructed base on the optimized SARS-CoV RBD-ACE2 complex structure using DS and was also optimized by Rosetta Relax. The interface between the RBD and ACE2 was analyzed by Rosetta Interface Analyzer^30^ and DS. Software PyMol was used for preparing structural figures^39^. Structural similarity analysis between SARS-CoV-2 and SARS-CoV was carried out by Discovery Studio (DS)^37^.

### Structural conservation analysis of SARS-CoV RBD

Structural conservation analysis of SARS-CoV RBD was carried out using ConSurf^35^. Alignment of 111 SARS-CoV RBD sequence, used for the generation of sequence conservation, was collected by BLAST via NCBI website^34^. Structural conservation was displayed by PyMol^39^.

## Declaration of interest statement

## Supplementary Material

**Table S1.** Interface analysis of RBD-ACE2 complex of SARS -CoV and SARS-CoV-2.

|  | **dG_separated**<br>**(REU)** | **dSASA_int**<br>**(Å<sup>2</sup>)** | **delta_unsat**<br>**Hbonds** | **hbonds_int** | **packstat** | **fa_atr**<br>**(REU)** | **fa_rep**<br>**(REU)** | **fa_elec**<br>**(REU)** |
| --- | --- | --- | --- | --- | --- | --- | --- | --- |
| **SARS-CoV** | -40.42 | 1838 | 7 | 10 | 0.621 | -4893 | 641.9 | -1193 |
| **SARS-CoV-2** | -44.96 | 1873 | 6 | 10 | 0.608 | -4944 | 654.8 | -1187 |

dG\_separated: The binding freeenergy. The change in Rosetta energy when the interface forming chains are separated, versus when they are complexed.
dSASA\_int: The solvent accessible area buried at the interface.
delta\_unsatHbonds: The number of buried, unsatisfied hydrogen bonds at the interface.
hbonds\_int: Total cross-interface hydrogen bonds found.
packstat: Rosetta's packing statistic score for the interface (0=bad, 1=perfect).
fa\_atr: Lennard-Jones attractive between atoms in different residues.
fa\_rep: Lennard-Jones repulsive between atoms in different residues.
fa\_elec: Coulombic electrostatic potential with a distance-dependent dielectric.

**Table S2.** RBD side chain interaction in the RBD-ACE2 interface.

| SARS-CoV |  | SARS-CoV-2 |  | Similarity | SARS-CoV |  |  |  |  | SARS-CoV-2 |  |  |  |  |
| --- | --- | --- | --- | --- | --- | --- | --- | --- | --- | --- | --- | --- | --- | --- |
| ID | Type | ID | Type |  | Distance | SB | HB | PI | NP | Distance | SB | HB | PI | NP |
| 390 | K | 403 | R | 67% | 6.5 | E37 |  |  |  | 4.6 | E37 |  |  |  |
| 392 | D | 405 | D | 100% | 6.4 | R393 |  |  |  | 6.5 | R393 |  |  |  |
| 404 | V | 417 | K | 24% | 7.4 |  |  |  |  | 7.6 | D30 |  |  |  |
| 426 | R | 439 | N | 42% | 4.0 | E329 |  |  |  | 7.4 |  |  |  |  |
| 432 | S | 445 | V | 25% | 6.6 |  |  |  |  | 5.6 |  |  |  | L45 |
| 436 | Y | 449 | Y | 100% | 2.9 |  | D38 | D38 |  | 2.8 |  | D38 | D38 |  |
| 440 | Y | 453 | Y | 100% | 3.3 |  |  | H34 |  | 3.0 |  |  | H34 |  |
| 442 | Y | 455 | L | 32% | 3.2 |  |  | H34 | K31 | 4.3 |  |  |  | H34,K31 |
| 443 | L | 456 | F | 45% | 3.7 |  |  |  | T27,K31 | 3.2 |  |  |  | T27,K31 |
| 460 | F | 473 | Y | 67% | 3.7 |  |  |  | T27 | 3.6 |  |  |  | T27 |
| 462 | P | 475 | A | 32% | 3.5 |  |  |  | Q24,T27 | 3.7 |  |  |  | Q24,T27 |
| 463 | D | 476 | G | 30% | 2.7 |  | S19 |  |  | 4.9 |  |  |  |  |
| 470 | P | 484 | E | 30% | 5.9 |  |  |  |  | 2.8 | K31 | K31 |  |  |
| 472 | L | 486 | F | 45% | 3.8 |  |  |  | M82 | 3.4 |  |  | Y83 | M82 |
| 473 | N | 487 | N | 100% | 2.7 |  | Q24,Y83 |  |  | 2.7 |  | Q24,Y83 |  |  |
| 475 | Y | 489 | Y | 100% | 2.8 |  | Y83 |  | T27,K31 | 2.8 |  | Y83 |  | T27,K31 |
| 479 | N | 493 | Q | 42% | 3.5 |  |  |  | H34 | 3.6 |  |  |  | H34 |
| 484 | Y | 498 | Q | 30% | 3.0 |  | Q42 | Y41 |  | 3.8 |  |  |  | Y41 |
| 486 | T | 500 | T | 100% | 2.7 |  | Y41 |  |  | 2.4 |  | Y41 |  |  |
| 487 | T | 501 | N | 42% | 3.5 |  |  |  | Y41,K353 | 3.3 |  |  |  |  |
| 489 | I | 503 | V | 88% | 4.6 |  |  |  | T324 | 4.3 |  |  |  | T324 |
| 491 | Y | 505 | Y | 100% | 3.0 |  | R393 |  | K353 | 2.9 |  | R393 |  | K353 |

Distance: the nearest distance from side chain of RBD residue to ACE2.
SB: salt-bridge. HB: hydrogen bond. PI: π-stack or π-anion interaction. NP: non-polar interaction.

**Figure S1.**
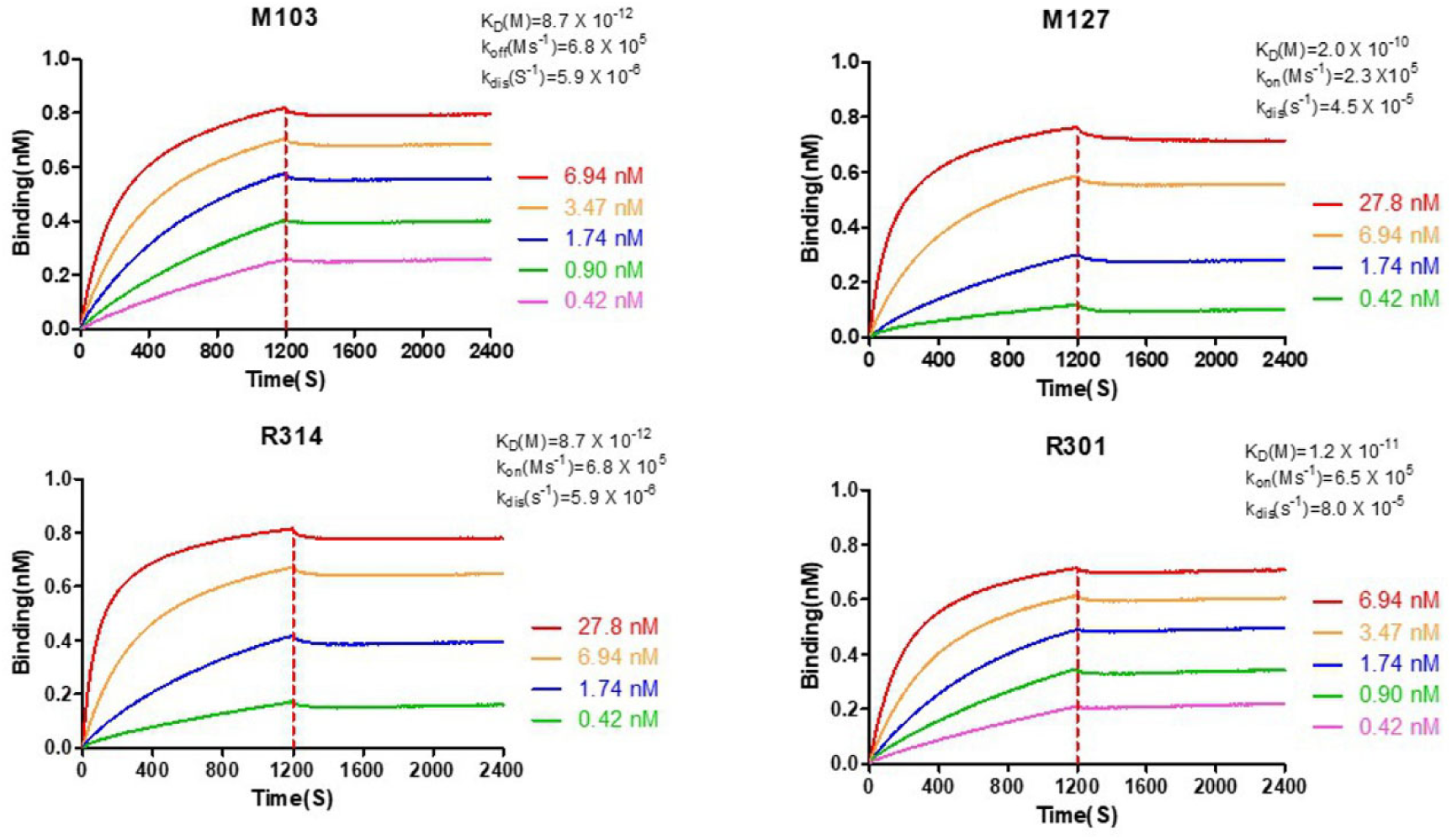
Realtime association and dissociation of monoclonal nAbs detected by Octet system.

**Figure S2.**
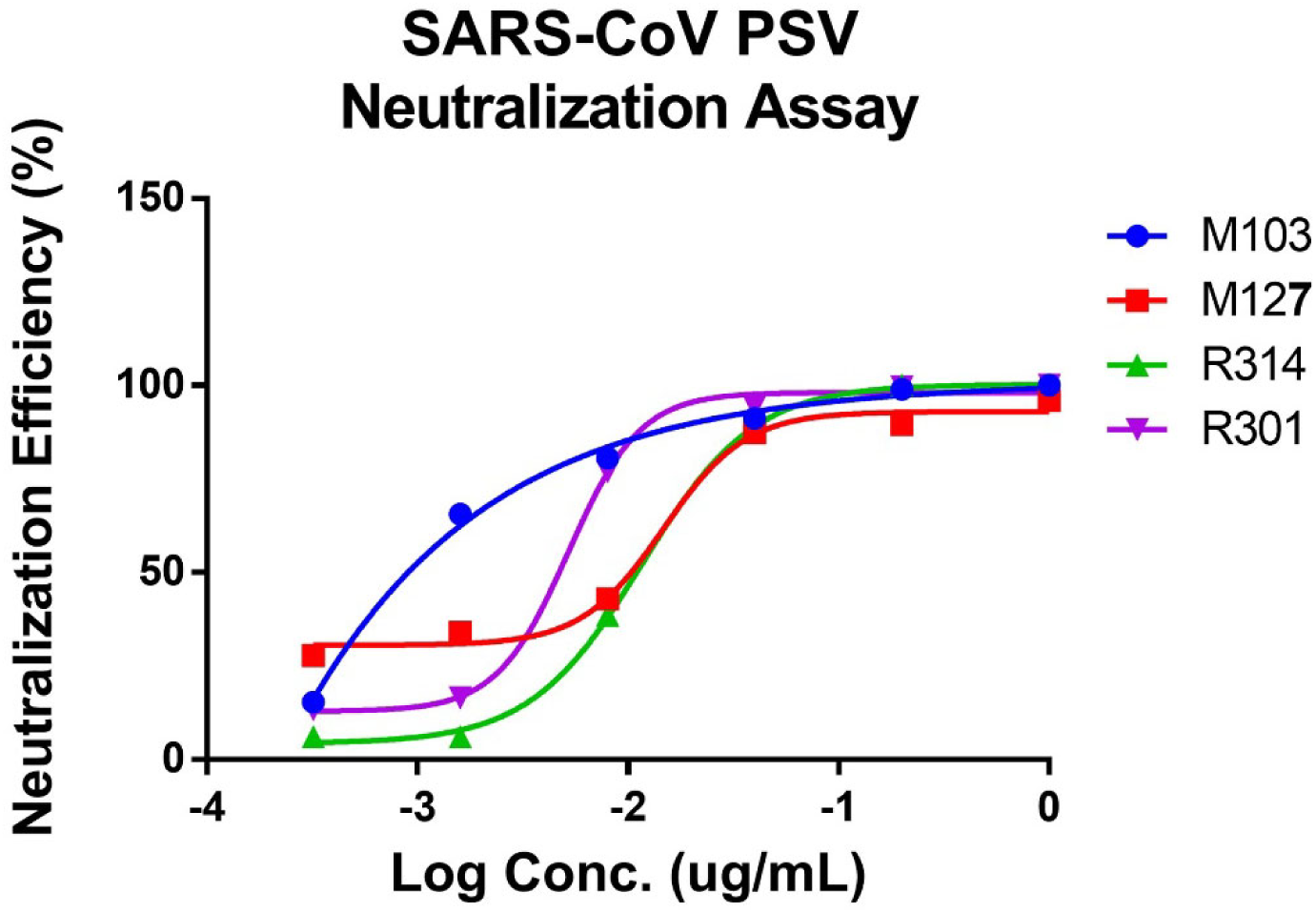
Neutralization assays of monoclonal nAbs against SARS-CoV PSV.

## References

1. Wu, P., et al. Real-time tentative assessment of the epidemiological characteristics of novel coronavirus infections in Wuhan, China, as at 22 January 2020. Eurosurveillance 25, 2000044 (2020).

2. Wang, C., Horby, P.W., Hayden, F.G. & Gao, G.F. A novel coronavirus outbreak of global health concern. Lancet (London, England) (2020).

3. Munster, V.J., Koopmans, M., van Doremalen, N., van Riel, D. & de Wit, E. A Novel Coronavirus Emerging in China - Key Questions for Impact Assessment. The New England journal of medicine (2020).

4. Ren, L.L., et al. Identification of a novel coronavirus causing severe pneumonia in human: a descriptive study. Chinese medical journal (2020).

5. The, L. Emerging understandings of 2019-nCoV. The Lancet395, 311 (2020).

6. Li, Q., et al. Early Transmission Dynamics in Wuhan, China, of Novel Coronavirus–Infected Pneumonia. New England Journal of Medicine (2020).

7. CDC. http://2019ncov.chinacdc.cn/nCoV/.

8. Yang, Y., et al. Epidemiological and clinical features of the 2019 novel coronavirus outbreak in China. medRxiv, 2020.2002.2010.20021675 (2020).

9. Tian, X., et al. Potent binding of 2019 novel coronavirus spike protein by a SARS coronavirus-specific human monoclonal antibody. bioRxiv, 2020.2001.2028.923011 (2020).

10. Lu, R., et al. Genomic characterisation and epidemiology of 2019 novel coronavirus: implications for virus origins and receptor binding. The Lancet (2020).

11. Fehr, A.R. & Perlman, S. Coronaviruses: an overview of their replication and pathogenesis. Methods in molecular biology (Clifton, N.J.)1282, 1–23 (2015).

12. Li, F. Receptor recognition mechanisms of coronaviruses: a decade of structural studies. Journal of virology 89, 1954–1964 (2015).

13. Song, Z., et al. From SARS to MERS, Thrusting Coronaviruses into the Spotlight. Viruses 11(2019).

14. Kuba, K., et al. A crucial role of angiotensin converting enzyme 2 (ACE2) in SARS coronavirus–induced lung injury. Nature Medicine 11, 875–879 (2005).

15. Coughlin, M., et al. Generation and characterization of human monoclonal neutralizing antibodies with distinct binding and sequence features against SARS coronavirus using XenoMouse. Virology 361, 93–102 (2007).

16. Li, F., Li, W., Farzan, M. & Harrison, S.C. Structure of SARS coronavirus spike receptor-binding domain complexed with receptor. Science 309, 1864–1868 (2005).

17. Wan, Y., Shang, J., Graham, R., Baric, R.S. & Li, F. Receptor recognition by novel coronavirus from Wuhan: An analysis based on decade-long structural studies of SARS. JVI.00127-00120 (2020).

18. Xu, X., et al. Evolution of the novel coronavirus from the ongoing Wuhan outbreak and modeling of its spike protein for risk of human transmission %J SCIENCE CHINA Life Sciences.

19. Walls, A.C., et al. Unexpected Receptor Functional Mimicry Elucidates Activation of Coronavirus Fusion. Cell1 76, 1026-1039.e1015 (2019).

20. Xu, J., et al. Antibodies and vaccines against Middle East respiratory syndrome coronavirus. 8, 841–856 (2019).

21. Pascal, K.E., et al. Pre- and postexposure efficacy of fully human antibodies against Spike protein in a novel humanized mouse model of MERS-CoV infection. Proceedings of the National Academy of Sciences 112, 8738–8743 (2015).

22. Zhu, Z., et al. Potent cross-reactive neutralization of SARS coronavirus isolates by human monoclonal antibodies. Proc Natl Acad Sci U S A 104, 12123–12128 (2007).

23. Greenough, T.C., et al. Development and characterization of a severe acute respiratory syndrome-associated coronavirus-neutralizing human monoclonal antibody that provides effective immunoprophylaxis in mice. J Infect Dis 191, 507–514 (2005).

24. ter Meulen, J., et al. Human monoclonal antibody as prophylaxis for SARS coronavirus infection in ferrets. Lancet (London, England) 363, 2139–2141 (2004).

25. Roberts, A., et al. Therapy with a severe acute respiratory syndrome-associated coronavirus-neutralizing human monoclonal antibody reduces disease severity and viral burden in golden Syrian hamsters. J Infect Dis 193, 685–692 (2006).

26. Sui, J., et al. Evaluation of human monoclonal antibody 80R for immunoprophylaxis of severe acute respiratory syndrome by an animal study, epitope mapping, and analysis of spike variants. Journal of virology 79, 5900–5906 (2005).

27. Rockx, B., et al. Escape from human monoclonal antibody neutralization affects in vitro and in vivo fitness of severe acute respiratory syndrome coronavirus. J Infect Dis 201, 946–955 (2010).

28. Tang, X., et al. Differential stepwise evolution of SARS coronavirus functional proteins in different host species. BMC evolutionary biology 9, 52–52 (2009).

29. Holmes, E.C. & Rambaut, A. Viral evolution and the emergence of SARS coronavirus. Philosophical transactions of the Royal Society of London. Series B, Biological sciences 359, 1059–1065 (2004).

30. Stranges, P.B. & Kuhlman, B. A comparison of successful and failed protein interface designs highlights the challenges of designing buried hydrogen bonds. Protein Sci 22, 74–82 (2013).

31. Du, L., et al. The spike protein of SARS-CoV--a target for vaccine and therapeutic development. Nat Rev Microbiol 7, 226–236 (2009).

32. Chakraborti, S., Prabakaran, P., Xiao, X. & Dimitrov, D.S. The SARS coronavirus S glycoprotein receptor binding domain: fine mapping and functional characterization. Virology journal 2, 73 (2005).

33. Gupta, R., Jung, E. & Brunak, S. Prediction of N-glycosylation sites in human proteins. 46, 203–206 (2004).

34. Altschul, S.F., Gish, W., Miller, W., Myers, E.W. & Lipman, D.J. Basic local alignment search tool. Journal of molecular biology 215, 403–410 (1990).

35. Ashkenazy, H., et al. ConSurf 2016: an improved methodology to estimate and visualize evolutionary conservation in macromolecules. Nucleic Acids Res 44, W344–350 (2016).

36. Sievers, F., et al. Fast, scalable generation of high-quality protein multiple sequence alignments using Clustal Omega. Molecular systems biology 7(2011).

37. Discovery Studio version 2.5. Accelrys Inc.: San Diego, CA, USA (2009).

38. Nivon, L.G., Moretti, R. & Baker, D. A Pareto-optimal refinement method for protein design scaffolds. PLoS One 8, e59004 (2013).

39. DeLano, W.L. The PyMOL molecular graphics system. http://www.pymol.org (2002).

